# The effect of long-term sleep disruption on the brain – looking beyond amyloid

**DOI:** 10.1101/2023.12.15.571841

**Authors:** L.J. Mentink, M.J.P. van Osch, L.J. Bakker, M.G.M. Olde Rikkert, C.F. Beckmann, J.A.H.R. Claassen, K.V. Haak

**Author notes:** Corresponding author Lara J. Mentink Department of Geriatrics, Radboud university medical center. These authors contributed equally as co-last authors (J.A.H.R. Claassen, K.V. Haak).

## Abstract

The mechanism underlying the possible causal association between long-term sleep disruption and Alzheimer’s disease remains unclear [1]. A hypothesised pathway through increased brain amyloid load was not confirmed in previous work in our cohort of maritime pilots with long-term work-related sleep disruption [2]. Here, using functional MRI, T2-FLAIR and Arterial Spin Labeling MRI scans, we explored alternative neuroimaging biomarkers related to both sleep disruption and AD: resting-state network co-activation and between-network connectivity of the default mode network (DMN), salience network (SAL) and frontoparietal network (FPN), vascular damage and cerebral blood flow (CBF).

We acquired data of 16 maritime pilots (56 ± 2.3 years old) with work-related long-term sleep disruption (23 ± 4.8 working years) and 16 healthy controls (59 ± 3.3 years old), with normal sleep patterns (Pittsburgh Sleep Quality Index ≤ 5). Maritime pilots did not show altered co-activation in either the DMN, FPN, or SAL and no differences in between-network connectivity. We did not detect increased markers of vascular damage in maritime pilots, and additionally, maritime pilots did not show altered CBF-patterns compared to healthy controls.

In summary, maritime pilots with long-term sleep disruption did not show neuroimaging markers indicative of preclinical AD compared to healthy controls. These findings do not resemble those of short-term sleep deprivation studies. This could be due to resiliency to sleep disruption or selection bias, as participants have already been exposed to and were able to deal with sleep disruption for multiple years, or to compensatory mechanisms [3]. This suggests the relationship between sleep disruption and AD is not as strong as previously implied in studies on short-term sleep deprivation, which would be beneficial for all shift workers suffering from work-related sleep disruptions.

## 1. Introduction

Sleep disturbances in midlife have been associated with an increased risk of late-onset Alzheimer’s disease (AD), yet the mechanism linking sleep to AD remains unclear [4–8]. Moreover, the relationship between AD and sleep is thought to be bidirectional [9]. Sleep-wake disturbances are a common symptom of moderate AD [10], but were found to already emerge in the preclinical stages of the disease [11]. Therefore, the association between sleep deprivation and AD could be due to causation as well as reverse causation.

Accumulation of the amyloid beta protein (Aβ) in the brain, a pathological hallmark associated with AD, has been hypothesized to be a possible underlying mechanism [9, 12]. In view of the circadian rhythm of Aβ production and clearance, increased levels of Aβ after sleep deprivation can be explained in two ways. Firstly, increased production of Aβ could be due to higher neuronal and synaptic activity during prolonged wakefulness [1, 9, 13]. Secondly, Aβ may accumulate as a result of decreased clearance of Aβ due to reduced total (deep) sleep time, as the glymphatic system for clearing toxins from the central nervous system is mainly active during sleep [14–16]. In support of this hypothesis, early animal studies indicated an increase in cortical Aβ deposition after sleep deprivation [17–20]. This was translated to humans, where the overnight reduction in Aβ levels in cerebrospinal fluid (CSF) after one night of normal sleep was absent after one night of full sleep deprivation. This was confirmed after one night of selective slow wave sleep disruption [21, 22]. In addition, Winer et al. (2020) observed that reduced slow wave sleep and sleep efficiency were associated with the rate of cortical Aβ accumulation [23]. Taken together, these studies provide some evidence for a causal link between sleep deprivation and AD via the Aβ cascade hypothesis. However, these studies remain limited to short-term effects of sleep on Aβ, and information on longer term effects or on development of AD is lacking.

In our “Sleep-Cognition Hypothesis In Maritime Pilots (SCHIP)” study, in which we investigated a unique group of older maritime pilots, no evidence for increased cortical Aβ accumulation has been found after long-term (approximately 20 years) intermittent sleep disruption, with reduced and fragmented sleep, related to their highly irregular shift work [2]. This finding was unexpected, in view of our hypothesis that the short-term increases in Aβ after sleep disruption, when repeated many times over the years, would result in chronic cortical Aβ accumulation [21, 22]. This therefore challenges the hypothesis that increased Aβ accumulation is the underlying cause of the association between sleep disruption and the increased risk for AD. This is in line with a recent study in mice where chronic sleep deprivation activated metabolic processes in the brain, independent of Aβ accumulation [24]. Therefore, in order to advance our understanding of associations between sleep and neuroimaging markers, we here look more broadly across sets of imaging-derived phenotypes and their association with sleep disruption.

Decreased co-activation of the default mode network (DMN) in a population of older adults is a biomarker of Alzheimer’s disease that might manifest prior to or independent of possible amyloid deposition [25]. The DMN is one of the major resting state networks, showing synchronous activity during rest from spatially distinct brain regions, measurable with functional MRI (fMRI) [26, 27]. The co-activation of the DMN decreases throughout the aging process, with accelerated deactivation in mild cognitive impairment (MCI) and AD [28–30]. It encompasses also the first regions to show accumulation of Aβ plaques, whereas co-activation is already affected before the Aβ accumulation is measurable with PET [31]. Interestingly, sleep disturbances also directly influence the DMN; one night of total sleep deprivation has been shown to significantly decrease DMN connectivity in healthy, young participants [32]. While the DMN is the most investigated network in the existing literature, a few studies indicate that other resting-state networks may also be affected in sleep deprivation, aging, and Alzheimer’s disease, i.e. the salience (SAL) and frontoparietal (FPN) networks [33–36]. Therefore we will extend this study on DMN with an exploratory analysis of the functional co-activation of the SAL and FPN networks and their inter-network connectivity.

Besides resting-state fMRI biomarkers, sleep disruption is potentially also linked with AD through cardiovascular pathways. Although this relation between sleep disruption and AD has not been widely studied, there are indications that disturbed circadian rhythms, and short sleep duration, and sleep quality are associated with cerebral small vessel disease and have other cardiovascular consequences such as reduced cerebral perfusion and increased white matter hyperintensities, although the directionality of these associations is not always clear [37–41]. Moreover, perfusion changes have been found very early in the disease cascade of AD [42]. Therefore, an additional aim of our study was to investigate vascular damage and cerebral flood flow (CBF). By looking beyond the amyloid hypothesis, the SCHIP study allows us insight into the possible mechanisms with which sleep disruptions could lead to AD.

## 2. Methods

### 2.1 Participants

Maritime pilots participating in the SCHIP study [2], were approached to participate in this follow-up study. Their profession (median years in profession = 24 years) causes these participants to have a highly irregular work and sleep schedule, consisting of a 7-day workweek during which they are on call for 24 hours a day, followed by a week off, leading to externally induced sleep disruptions. For details on this unique population, see [2, 43]. From the original study population (n= 19), 16 maritime pilots (median age = 56 years, age range = 52-61 years) agreed to participate in this follow-up study (table 1). In addition, we recruited 16 healthy male controls participants matched for age (median age = 59 years, age range = 51 – 66 years), education level and physical activity. The control participants did not perform shift work and had normal sleep, confirmed by a Pittsburgh Sleep Quality Index of ≤ 5 for the previous month, and regular sleep/wake schedules. Exclusion criteria were: use of psychoactive medication, consuming > 30 alcoholic beverages per week, BMI > 30 kg/m^2^, diagnosed with an intrinsic sleeping disorder. The SCHIP study was approved by the institutional review board (IRB) (CMO Region Arnhem-Nijmegen, NL55712.091.16; file number 2016-2337) and was performed in accordance with Good Clinical Practice (GCP) guidelines. We obtained written informed consent from all participants.

**Table 1:** Baseline characteristics of the maritime pilots participating in this SCHIP study follow-up.

|  | Maritime pilots (N=16) | Healthy controls (N=16) | p-value |
| --- | --- | --- | --- |
| **Age, years** | 56 (54 – 59) | 59 (57 – 60.5) | **0.04** |
| **Sex, males (%)** | 16 (100) | 16 (100) | 1 |
| **Education, ISCED score** | 7 (7 - 7) | 6.5 (6 - 7) | **0.008** |
| **BMI, kg/m<sup>2</sup>** | 26.6 (22.7 – 27.0) | 25.3 (23.7 – 28.1) | 0.95 |
| **Working years, maritime pilots only** | 24 (19.5 – 27) | - | - |
| **History of high cholesterol, n(%)** | 1 (6) | 1 (6) | 1 |
| **History of high blood pressure, n(%)** | 1 (6) | 2 (13) | 0.58 |
| **Physical activity > 30 minutes a day, n(%)** | 14 (88) | 15 (94) | 0.58 |
| *once a week, n(%)* | 2 (13) | 1 (6) | 0.58 |
| *twice a week, n(%)* | 4 (25) | 2 (13) | 0.39 |
| *≥ three times a week, n(%)* | 8 (50) | 12 (75) | 0.16 |
| **Medication use, n(%)** | 3 (19) | 5 (31) | 0.44 |
| *antihypertensives, n(%)* | 0 (0) | 2 (13) | 0.16 |
| *statins, n(%)* | 0 (0) | 1 (6) | 0.35 |
| *other, n(%)* | 3 (19) | 4 (25) | 0.69 |
| **Smoking, n(%)** | 1 (6) | 2 (13) | 0.58 |
| **Alcohol use, number of drinks per week** | 3 (2 – 6.5) | 2 (1 – 5.5) | 0.38 |
| **Hours since last shift, non-retired maritime pilots only (N=11)** | 73.5 (69 - 143) | - | - |

Notes: Data displayed as median (IQR) unless otherwise stated. Abbreviations: PSQI, Pittsburgh Sleep Quality Index; BMI, Body Mass Index; ISCED, International Standard Classification of Education; SD, standard deviation; IQR, interquartile range. Group differences were tested using a Mann-Whitney U test, for non-normally distributed data.

### 2.2 Neuroimaging protocol & data pre-processing

The participants were scanned on a 3T MR scanner (Prisma, Siemens, Germany) at the Donders Centre for Cognitive Neuroimaging. The MR protocol entailed the acquisition of structural T1-weighted MRI, resting-state fMRI, T2-weighted FLAIR and Arterial Spin Labeling (ASL) perfusion scans.

#### 2.2.1 Resting-state fMRI

Resting-state data were acquired using a 10-minute multiband accelerated EPI sequence (TR = 1000 ms, TE = 34 ms, flip angle = 60 degrees, spatial resolution = 2.0 mm isotropic, number of slices = 66, multiband factor = 6). Additionally, we acquired spin-echo EPI data with reversed phase-encode blips for distortion correction purposes (TR =5186 ms, TE = 47.40 ms, flip angle = 90 degrees, spatial resolution = 2.0 mm isotropic).

Fieldmaps were created using FSL’s topup; Data was collected with reversed phase-encode blips, resulting in pairs of images with distortions going in opposite directions. From these pairs the susceptibility-induced off-resonance field was estimated using a method similar to that described in [44] as implemented in FSL [45] and the two images were combined into a single corrected one.

Preprocessing of the resting-state fMRI data was performed in FSL’s FEAT. This included removal of the first five volumes, fieldmap correction, realignment to the middle volume using MCFLIRT [46], global intensity normalisation, and 6mm FWHM spatial smoothing. We then used ICA-AROMA, a data-driven ICA-based method to remove signal components related to head motion [47]. Afterwards, nuisance regression was performed to the white matter and cerebrospinal fluid (CSF) signal. Lastly, we applied a highpass temporal filter with cut-off frequency of 0.01Hz and registered the data to 2mm MNI152 space.

#### 2.2.2 FLAIR

Data were acquired using a 3D FLAIR sequence with an acquisition time of 5 minutes 12 seconds (TR = 5000 ms, TE = 397 ms, TI = 1800 ms, spatial resolution = 0.5 × 0.5 × 1 mm, GRAPPA acceleration factor = 3).

The MRI T2-FLAIR scans were used to visually score periventricular lesions (PVL), white matter hyperintensities (WMH), lacunes (LAC) and perivascular spaces (PVS). Assessors (LM & LB) were first trained in scoring by an experienced assessor (JC). Sufficient training was confirmed by JC after scoring a practice dataset with T2-FLAIR scans collected at the department of Geriatrics. For scoring the severity of WMH, the Fazekas scale [48] was used. The Fazekas scale makes a distinction between PVL and DWMH, and ranges between 0 and 3. For PVL, 1 is scored in case of caps and a pencil-thin lining (< 5 mm), 2 is scored in case of a smooth halo (> 5 mm), and 3 is scored in case of an irregular cap extending into deep white matter. For DWMH, 0 was scored in case of no MRI-visible DWMH, 1 in case of single lesion of ≤ 9 mm or grouped lesions of < 20 mm, 2 in case of single lesions between 10 and 20 mm or grouped lesions of > 20 mm with no more than connecting bridges between individual lesions, and 3 was scored in case of single lesions or confluent areas of ≥ 20 mm. The number of lacunes (> 3 mm) was counted and enlarged PVS were assessed using a semi-quantitative scale ranging from 0 up to 4 whereby a score 0 is given in case of no MRI-visible PVS, 1 in case of <10 PVS, 2 in case of 10 to 20 PVS, 3 in case of 21 to 40 PVS and 4 in case of > 40 PVS [49, 50]. Table 2 lists the cut-off points to score PVL, DWMH and PVS. In case of disagreement in scoring between assessors, a third researcher (JC) would score the item.

**Table 2:** Scoring matrix for periventricular lesions, deep white matter hyperintensities, and perivascular spaces.

| Score → | 0 | 1 | 2 | 3 | 4 |
| --- | --- | --- | --- | --- | --- |
| Item ↓ |  |  |  |  |  |
| PVL | No PVL | Caps and/or pencil-thin lining ($< 5$mm) | Smooth halo ($> 5$mm) | Irregular caps extending into deep white matter | - |
| DWMH | No DWMH | Single lesions $\leq 9$mm or grouped lesions of $\leq 20$mm | Single lesions $\geq 10$mm and $< 20$mm or grouped lesions $> 20$mm with no more than connecting bridges between individual lesions | Single lesions or confluent areas of $\geq 20$mm | - |
| PVS | No PVS | <10 | 10-20 | 21-40 | >40 |

Notes: PVL, periventricular lesions; DWMH, deep white matter hyperintensities; PVS, perivascular spaces.

#### 2.2.3 Arterial Spin Labeling MRI

ASL data were acquired using the Siemens standard pulsed ASL FAIR-QUIPSSII sequence (6 measurements, single-TI of 1990 ms, bolus duration = 700 ms), with a 3D readout (TR = 4600 ms, TE = 16.28 ms, spatial resolution = 3 mm isotropic, interpolated to 1.5 x 1.5 x 3 mm, 40 slices, echo planar imaging factor = 21).

With FSL’s MCFLIRT we registered the ASL measurements to the middle volume. Using FSL’s aslfile we then calculated the difference data by subtracting the tag-control pairs, as well as calculated a mean control image to use as a substitute for the M0 image. To calculate a measure of cerebral blood flow (CBF), we used the formula recommended by the ISMRM perfusion study group and the European consortium for ASL in dementia (formula 1 and 2) [51] , but due to the absence of a proper M0, further analysis focuses on relative changes in CBF by including the participant’s average whole brain GM CBF values as covariate in the statistical analysis.

To determine region-specific CBF-values, we selected the hippocampi and posterior cingulate cortex (PCC) from the HarvardOxford cortical and subcortical 2mm atlas and registered these ROIs from standard to native space with FSL’s FLIRT (FMRIB’s Linear Image Registration Tool) and FNIRT (FMRIB’s Linear Image Registration Tool) [46, 52]. We segmented the T1 image with FSL’s FAST and then similarly to above registered the segmented grey matter to native space [53]. By combining the binarized ROI’s with the binarized grey matter mask, we selected only the grey matter voxels within our ROI’s. We then utilized FSL’s fslstats to calculate the average value of CBF of the grey matter inside these ROI’s. Additionally, we calculated the average value of CBF in the whole-brain grey matter.

For the voxel-wise analysis, we dilated the whole-brain mask output from FAST and registered this mask to native space to create a brain-extracted CBF image. We then registered the CBF images to standard space (MNI152) using FLIRT and FNIRT [46, 52].

### 2.3 Statistical analyses

#### 2.3.1 Resting-state fMRI

A group-level independent component analysis, implemented in FSL’s MELODIC (Multivariate Exploratory Linear Optimised Decomposition into Independent Components) identified 25 resting-state networks [54]. Using dual regression, the subject-specific spatial maps and associated timeseries were estimated [55, 56]. The spatial map representing the DMN, SAL and FPN were selected based on visual inspection of the group ICA results.

The between-group differences were assessed using FSL’s Randomise permutation-testing tool (with TFCE inference, 10.000 permutations), with a Bonferroni-corrected p-value threshold of 0.05/6=0.008 as two contrasts were assessed in each of the three networks. Additionally, we added covariates of age, and, within the group of maritime pilots, history of work-years and, for non-retired maritime pilots, the duration between last shift and MRI scan (representing any short term sleep deprivation).

Four of the resulting 25 networks showed clear artefacts and were excluded from further analysis. As a quality check, we assessed the group means of all remaining ICA networks (21 of 25 networks), to indicate whether the mean response of each network is different from zero.

In addition to assessing the between-group co-activation of the three RSN’s, we were also interested if the correlation of the timeseries between the RSN’s differed between groups; i.e. whether there is a between-group difference in connection strength between the RSN’s. With FSLNets, we were able to calculate the full correlation matrix between the subject-specific timeseries of each RSN, the stage 1 dual regression output. Between-group differences were assessed using FSL’s Randomise with 10.000 permutations and corrected for the multiple comparisons across all edges.

#### 2.3.2 3D Flair

Normality in the lesion scores was tested with a Shapiro-wilk test. Between-group differences in vascular lesions were assessed using a Mann-Whitney-U test in MATLAB 2022b (Massachusetts, USA).

#### 2.3.3 Arterial Spin Labeling

The average CBF values in the ROIs were non-normally distributed. Therefore non-parametric permutation tests (FSL’s permutation analysis of linear models (PALM) with 10.000 permutations) were employed to assess group differences in CBF. Average CBF in whole-brain grey matter and summed probabilities of grey matter in the ROI’s for partial volume correction were added as a covariates to the model.

For the voxel-wise analyses, we also tested for group differences using non-parametric permutation tests (FSL’s Randomise, 10.000 permutations, with TFCE inference), this time using the whole brain CBF images in standard space as input. Average CBF in whole-brain grey matter was added as a covariate to the model.

## 3. Results

To study whether maritime pilots with long-term sleep disruption show altered brain biomarkers related to sleep and AD, we compared the 16 maritime pilots with 16 healthy matched controls.

### 3.1 Resting-state fMRI

For the main analyses, i.e. the resting-state fMRI, we studied the hypothesis that maritime pilots show altered RSN co-activation compared to controls; more specifically we hypothesized altered DMN, SAL and FPN co-activation. Using group-ICA and dual regression we were able to identify these networks in each subject and study between-group differences using permutation analyses. Our quality check indicated that the mean co-activation of the 21 meaningful networks from the 25 components was different from zero (p>0.0001 for all 21 networks in both groups) (figure 1).

**Figure 1:**
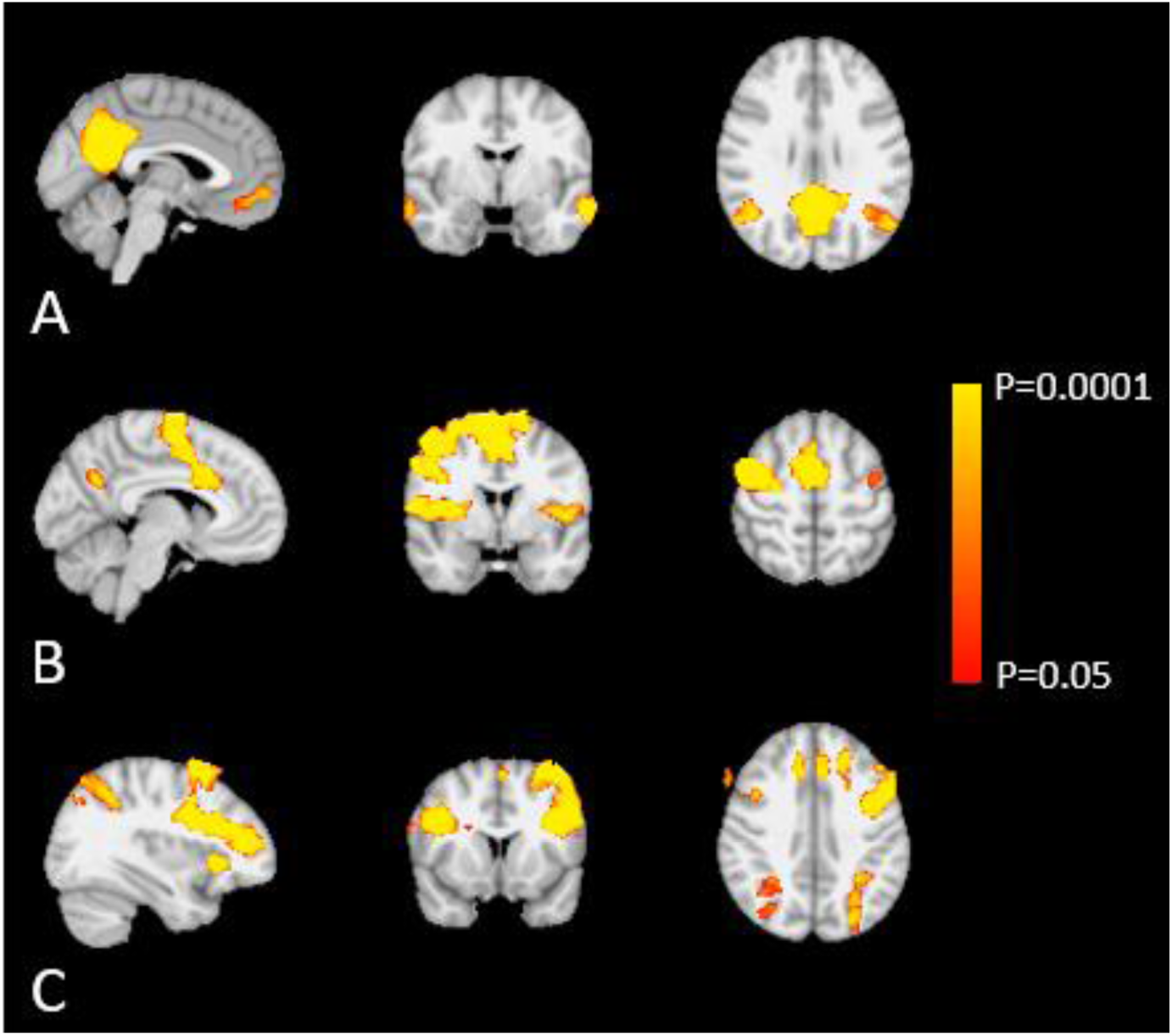
The mean co-activation of the default-mode (A), salience (B) and frontoparietal (C) networks.

No significant differences in DMN (pilots>controls: all p>0.28 and t<5.9; controls>pilots: all p>0.59 and t<5.0), SAL (pilots>controls: all p>0.24 and t<4.8; controls>pilots: all p>0.51 and t<4.9) and FPN (pilots>controls: all p>0.71 and t<4.9; controls>pilots: all p>0.46 and t<5.0) co-activation were detected. Including covariates of age, amount of work-years and short-term sleep deprivation did not alter these results.

Additionally, we were interested in whether there were between-group differences in the connection strength between the networks. Therefore, we tested the full correlation matrix using the subject-specific time courses of these networks. Our analysis did not reveal any significant differences in the intercorrelation of the three RSN’s between the maritime pilots and controls (DMN-SAL: uncorrected p>0.3; DMN-FPN: uncorrected p>0.21; SAL-FPN: uncorrected p>0.29; corrected p=1 for the connection strength between the three networks in both directions).

### 3.2 3D Flair

The intra-class correlation coefficient (ICC) showed moderate to good agreement between the two assessors (ICC=0.50 for PVS, ICC=0.79 for LAC, ICC=0.86 for DWMH, ICC=0.87 for PVL). Table 3 shows the scores for the maritime pilots and controls, displayed as median and interquartile range. The vascular scores do not differ between the two groups.

**Table 3:** Vascular scores for maritime pilots and healthy controls, displayed as median (IQR) and p-values of between-group differences in vascular scores..

| Item | Maritime pilot | Healthy control | p-value | Cohen's d |
| --- | --- | --- | --- | --- |
| PVL | 1 (0.5) | 1 (1) | 0.72 | 0.07 |
| DWMH | 1 (0) | 1 (0) | 0.77 | 0.07 |
| LAC | 0 (0) | 0 (0) | 1 | 0 |
| PVS | 2 (1) | 2 (0) | 0.44 | 0.14 |

Notes: PVL, periventricular lesions; DWMH, deep white matter hyperintensities; LAC, lacunes; PVS, perivascular spaces

### 3.3 Arterial Spin Labeling

We detected no significant differences in average CBF in the grey matter of the hippocampi and PCC ROIs, corrected for average CBF in whole-brain grey matter and summed probabilities of grey matter in the ROI’s for partial volume correction (hippocampus (pilots>controls: p=0.84; controls>pilots: p=0.16; d=0.18), PCC (pilots>controls: p=0.89; controls>pilots: p=0.11; d=0.12)) (figure 2).

**Figure 2:**
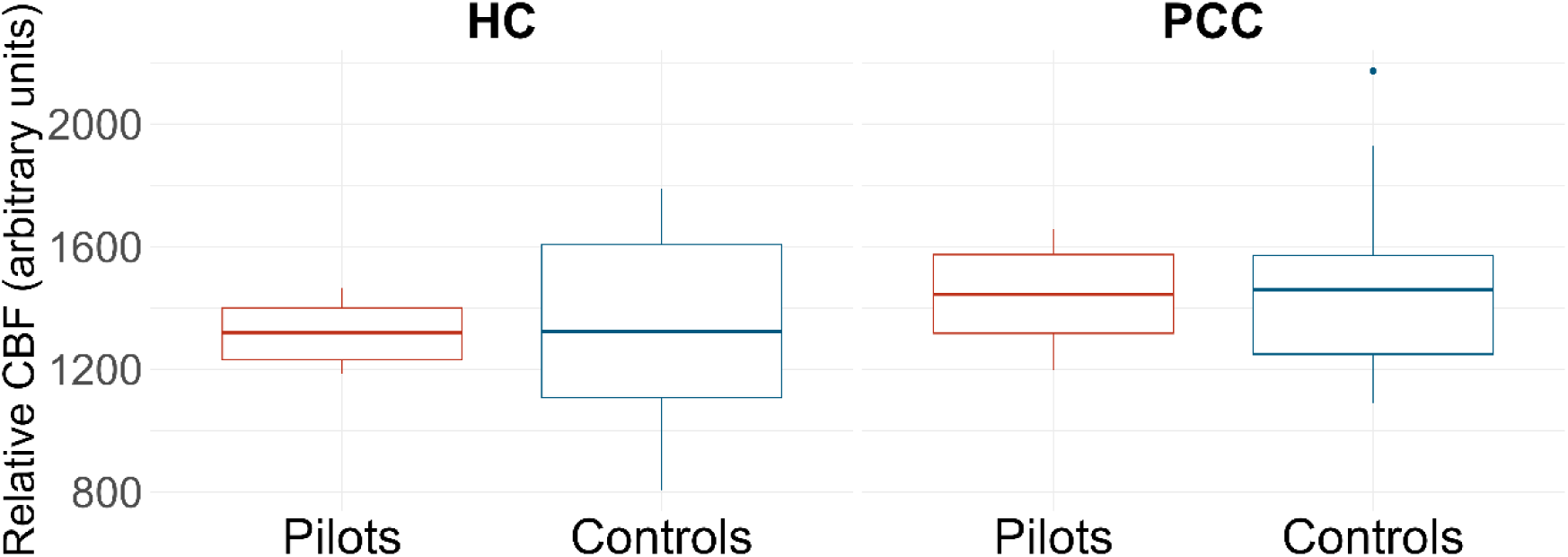
Boxplots indicating the spread in relative CBF in the hippocampi and PCC ROIs between pilots and controls. Abbreviations: CBF, cerebral blood flow; PCC, posterior cingulate cortex; ROI, region-of-interest.

In the whole-brain voxelwise analyses we found a marginally significant cluster of 3 voxels in the precuneus (corrected p>0.086; t<5.4) in the pilots>controls contrast (figure 3).

**Figure 3:**
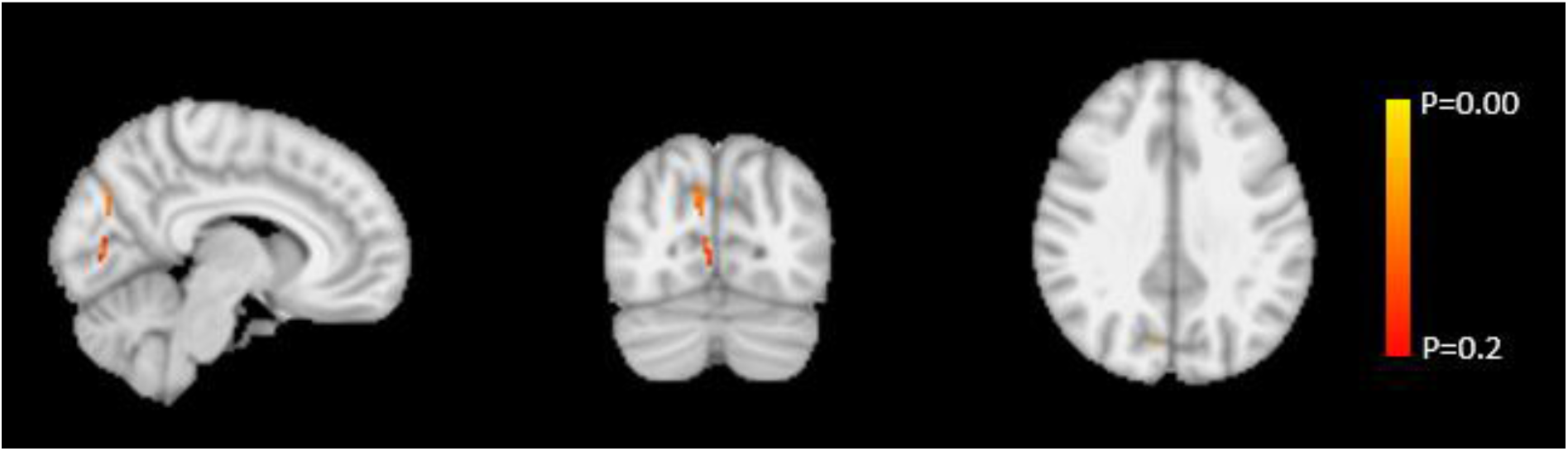
whole-brain voxelwise results, showing a small cluster of voxels in the precuneus with higher CBF in maritime pilots compared to controls (uncorrected p>0.043).

## 4. Discussion

In this study we were interested to explore neuroimaging biomarkers related to both sleep disruption and AD. In a previous study in these pilots [2] we found that long-term sleep disruption was not associated with Aβ accumulation, despite previous evidence from studies investigating the effects of short-term sleep deprivation [21, 22]. Therefore, we now aimed to explore other underlying mechanisms that could explain the epidemiological evidence linking long term sleep disruption to an increased risk for AD. We used a multimodal neuroimaging approach to study putative early brain changes known to be associated with sleep deprivation and/or increased risk for AD [33–36]. Specifically, we were interested in differences in resting-state network co-activation, vascular damage and cerebral perfusion between participants with long-term sleep disruption, i.e. maritime pilots, and healthy controls.

Using resting-state fMRI, we studied co-activation of the default mode network (DMN), salience network (SAL) and frontoparietal network (FPN). While the DMN is an established early biomarker for AD [25, 30], the SAL and FPN are rather understudied in the field of AD. However, these networks might be affected as a previous study has indicated the triple-network model consisting of DMN, SAL and FPN to be affected in AD and the same networks were affected after short-term sleep deprivation [57, 58]. Our results showed no differences between maritime pilots and healthy controls, in contrast to short-term sleep deprivation studies, where decreased DMN, SAL and FPN activations were found after acute total sleep deprivation [32, 36, 59]. While multiple studies are available on acute total sleep deprivation, evidence from long-term sleep disruption as assessed here is lacking. Therefore, we cannot be sure whether decreased RSN co-activation is a short-term effect, only occurring after acute total sleep deprivation, or whether it could still be an effect of long-term sleep disruption without any compensatory pauses. Additionally we were interested whether the connectivity strength between the three RNS’s differed between groups, however, no significant differences in between-network connectivity strength were found in our data between maritime pilots and controls. As this has not been studied in relation to sleep disruption, the between-network connectivity strength might only be affected in a later stage of AD, or in sleep disruption without compensatory pauses. We found no indication that it is an early biomarker of the disease.

A few studies have explored the relationship between sleep disruption and cerebrovascular damage, due to for example increased oxidative stress and blood-brain-barrier impairment [60]. This could be an alternative underlying mechanism between sleep disruption and AD [61, 62]. Results yielded in previous studies are mixed, showing both increased DWMH and increased small vessel disease in middle-aged and older adults with self-reported short sleep duration [40, 41, 63], as well as no association between short sleep duration at midlife and late life vascular damage and AD [64]. In line with Lutsey et al. [64], we found no association between sleep disruption and increased vascular damage, as vascular scores in both groups correspond to observations in healthy older adults [50]. However, direct comparison between studies is impeded due to the differences in sleep disruption between study populations (self-reported short sleep duration vs long-term work-related sleep disruptions). Two studies have investigated disrupted circadian rhythms, more closely resembling the sleep disruption of our study population. These studies both found an association between disturbed 24-hour activity rhythms and increased burden of cerebral small vessel disease, however, the directionality of this association could not be confirmed [37, 39]. A confounder in sleep disruption studies might be the (unknown) presence of obstructive sleep apnoea (OSA). Multiple studies in populations with OSA investigated its relationship with AD through vascular damage, because people with OSA have higher cardiovascular risk and has increased oxidative stress due to their nocturnal hypoxemia, possibly exacerbating neurovascular damage [64–67], as described in detail in the review by Daulatzai (2015) [68]. Indeed, the meta-analysis by Bubu et al. revealed that among sleep disorders, OSA carries the greatest relative risk (RR) for AD [4].

In our population of maritime pilots with long term sleep disruptions, we specifically explored CBF in the hippocampus and PCC, as regional CBF changes in areas already affected in early AD are thought to drive global CBF changes [69]. In addition we investigated whole brain grey and white matter CBF. We found no differences, compared to normal controls, in the hippocampal and PCC ROI’s in the maritime pilots. Our whole brain voxelwise analyses showed a small cluster with marginally increased CBF in maritime pilots (uncorrected p>0.043), however this did not survive correction for multiple testing. Comparison is limited due to the nature of the sleep disruption studied here, as previous work has been limited to acute short-term sleep deprivation experiments. Such studies found decreased cerebral perfusion after total sleep deprivation and sleep restriction in healthy young adults [70, 71].

Several hypotheses could explain our results. Firstly, there may be a true null effect in the causal link between externally-induced long-term sleep disruptions and the development of AD. Taking a closer look at the large meta-analysis by Bubu et al. (2016), the highest RR for AD was found in OSA, a sleep-related breathing disorder [4]. Furthermore, most of the included studies examined middle-aged and older individuals with a short follow-up. This could suggest that the associations could be driven by a specific sleep disorder (e.g. OSA) or by reverse-causality, wherein the reported sleep disorder is an early symptom of AD. An alternative interpretation of our results could be that a causal link between sleep disruptions and AD does exist, but was not detectable in our small sample size. However, our group had a median exposure of 24 years to sleep disruption. Therefore, if sleep disruption is indeed a relevant causal contributor to AD, such a long term and consistent exposure should be expected to show an effect, even in a relatively small sample, especially since we examined sensitive neuroimaging biomarkers of network systems involved in early stages of AD. Thirdly, there may be a causal link between poor sleep and AD, but compensation for sleep disruptions is possible as the maritime pilots have a week off after each workweek, which may have safeguarded them from long-term sleep deprivation. To avoid any confounding factors of intrinsic sleep disorders, we included a population with externally-induced sleep disruption. Our participants continued this occupation for a long time, which they might not have done when they would experience significant negative effects from shift work. We therefore may have selected a highly resilient population. Moreover, previous work in this group of maritime pilots illustrates a possible compensation mechanism of increasing total sleep time and the relative amount of deep sleep during their week off [3]. Fourthly, the causal link may be only present in a selected population. Recent research has demonstrated the multifactorial nature of AD, where the emergence of the disease is driven by multiple causal factors, therefore, sleep disruption in itself might not be enough to initiate AD development [72]. In contrast, our participants are healthy and, to our knowledge, have limited other risk factors for AD. An interesting factor to assess would have been APOE ε4-status, as APOE e4-carriership might modulate the relationship between sleep disruption and AD [73], however our ethical review board at the time did not allow individual APOE genotype determination in this study.

For future studies, we would suggest including a larger and more diverse population with regard to ancestry, sex, level of education, and lifestyle factors. When studying a population with work-related sleep disruption (i.e. shift work), it is important to carefully define their work and sleep schedules as well as the severity of sleep disruption and quantified (lack of) deep sleep, as research in the Danish Nurse Cohort has shown that not all types of shift work are associated with increased risk for AD [74], impacting generalizability across types of shift work and towards the general population. Furthermore, the field requires more longitudinal studies starting at a younger age compared to the studies in the large meta-analyses [4, 75], to accurately explore whether we are studying a causal relationship as opposed to reverse-causation.

In conclusion, we studied long-term work-related sleep disruption in an interesting group of shift-workers, maritime pilots. We decided to look beyond the amyloid hypothesis by investigating possible alternative underlying mechanisms affecting resting-state networks and cerebrovascular markers, which may present prior to or independent of Aβ accumulation. Here, we did not find evidence of impacted resting-state network co-activation, vascular damage or cerebral perfusion in our group of maritime pilots compared to healthy controls, which does not support a causal association between this type of sleep disruption and increased risk for AD. As the existing research predominantly concentrates on acute total sleep deprivation, which likely does not represent potential long-term risks, we believe that more longitudinal studies need to be conducted on long-term sleep disruptions to better understand the bidirectional relationship between sleep and AD, along with exploring potential compensation strategies.

## Acknowledgements

This work was supported by the Netherlands Organisation for Scientific Research (NWO) Vici Grant No. 17854 (to CB), NWO-CAS Grant No. 012-200-013 (to CB), Vici Grant No. 016.160.351 (to MvO), Vidi Grant No. 09150171910043 (to KH) and Veni Grant No. 016.Veni.171.068 (to KH) and Internationale Stichting Alzheimer Onderzoek (now Alzheimer Nederland) Grant No. 15040 (to JC), and a Wellcome Trust Collaborative Award 215573/Z/19/Z (to CB).

## Data availability statement

The data that support the findings of this study will be placed in a restricted access data sharing collection (DSC) on the Donders Repository and made available upon request.

## References

1. Musiek, E.S., D.D. Xiong, and D.M. Holtzman, Sleep, circadian rhythms, and the pathogenesis of Alzheimer disease. Experimental & molecular medicine, 2015. 47(3): p. e148.

2. Thomas, J., et al., Effects of long-term sleep disruption on cognitive function and brain amyloid-β burden: a case-control study. Alzheimer’s Research & Therapy, 2020. 12(1): p. 101.

3. Mentink, L.J., et al., Home-EEG assessment of possible compensatory mechanisms for sleep disruption in highly irregular shift workers – The ANCHOR study. PLOS ONE, 2021. 15(12): p. e0237622.

4. Bubu, O.M., et al., Sleep, Cognitive impairment, and Alzheimer’s disease: A Systematic Review and Meta-Analysis. Sleep, 2016. 40(1).

5. Musiek, E.S. and Y.-E.S. Ju, Targeting Sleep and Circadian Function in the Prevention of Alzheimer Disease. JAMA Neurology, 2022.

6. Sabia, S., et al., Association of sleep duration in middle and old age with incidence of dementia. Nature Communications, 2021. 12(1): p. 2289.

7. Lutsey, P.L., et al., Sleep characteristics and risk of dementia and Alzheimer’s disease: The Atherosclerosis Risk in Communities Study. Alzheimer’s & Dementia, 2018. 14(2): p. 157–166.

8. Luojus, M.K., et al., Self-reported sleep disturbance and incidence of dementia in ageing men. Journal of Epidemiology and Community Health, 2017. 71(4): p. 329–335.

9. Ju, Y.-E.S., B.P. Lucey, and D.M. Holtzman, Sleep and Alzheimer disease pathology--a bidirectional relationship. Nature reviews. Neurology, 2014. 10(2): p. 115–119.

10. Moran, M., et al., Sleep disturbance in mild to moderate Alzheimer’s disease. Sleep Medicine, 2005. 6(4): p. 347–352.

11. Ju, Y.-E.S., et al., Sleep quality and preclinical Alzheimer disease. JAMA neurology, 2013. 70(5): p. 587–593.

12. Lucey, B.P., et al., Effect of sleep on overnight cerebrospinal fluid amyloid β kinetics. Annals of Neurology, 2018. 83(1): p. 197–204.

13. Cirrito, J.R., et al., Synaptic Activity Regulates Interstitial Fluid Amyloid-β Levels In Vivo. Neuron, 2005. 48(6): p. 913–922.

14. Jessen, N.A., et al., The Glymphatic System: A Beginner’s Guide. Neurochem Res, 2015. 40(12): p. 2583–99.

15. Mestre, H., Y. Mori, and M. Nedergaard, The Brain’s Glymphatic System: Current Controversies. Trends in Neurosciences, 2020. 43(7): p. 458–466.

16. Xie, L., et al., Sleep Drives Metabolite Clearance from the Adult Brain. Science, 2013. 342(6156): p. 373–377.

17. Kang, J.-E., et al., Amyloid-β dynamics are regulated by orexin and the sleep-wake cycle.Science, 2009. 326(5955): p. 1005–1007.

18. Minakawa, E.N., et al., Chronic sleep fragmentation exacerbates amyloid β deposition in Alzheimer’s disease model mice. Neuroscience letters, 2017. 653: p. 362–369.

19. Rothman, S.M., et al., Chronic mild sleep restriction accentuates contextual memory impairments, and accumulations of cortical Aβ and pTau in a mouse model of Alzheimer’s disease. Brain research, 2013. 1529: p. 200–208.

20. Zhao, H.Y., et al., Chronic Sleep Restriction Induces Cognitive Deficits and Cortical Beta-Amyloid Deposition in Mice via BACE 1-Antisense Activation. CNS neuroscience & therapeutics, 2017. 23(3): p. 233–240.

21. Ju, Y.-E.S., et al., Slow wave sleep disruption increases cerebrospinal fluid amyloid-β levels. Brain, 2017. 140(8): p. 2104–2111.

22. Ooms, S., et al., Effect of 1 night of total sleep deprivation on cerebrospinal fluid β-amyloid 42 in healthy middle-aged men: a randomized clinical trial. JAMA neurology, 2014. 71(8): p. 971–977.

23. Winer, J.R., et al., Sleep Disturbance Forecasts β-Amyloid Accumulation across Subsequent Years. Current Biology, 2020. 30(21): p. 4291–4298.e3.

24. Parhizkar, S., et al., Sleep deprivation exacerbates microglial reactivity and Aβ deposition in a TREM2-dependent manner in mice. 2023. 15(693): p. eade6285.

25. Jack, C.R., et al., Tracking pathophysiological processes in Alzheimer’s disease: an updated hypothetical model of dynamic biomarkers. The Lancet Neurology, 2013. 12(2): p. 207–216.

26. Beckmann, C.F., et al., Investigations into resting-state connectivity using independent component analysis. Philosophical Transactions of the Royal Society B: Biological Sciences, 2005. 360(1457): p. 1001-13.

27. Raichle, M.E., et al., A default mode of brain function. Proceedings of the National Academy of Sciences, 2001. 98(2): p. 676–682.

28. Hafkemeijer, A., J. van der Grond, and S.A.R.B. Rombouts, Imaging the default mode network in aging and dementia. Biochimica et Biophysica Acta (BBA) - Molecular Basis of Disease, 2012. 1822(3): p. 431–441.

29. Damoiseaux, J.S., et al., Reduced resting-state brain activity in the “default network” in normal aging. Cerebral cortex, 2007. 18(8): p. 1856–1864.

30. Greicius, M.D., et al., Default-mode network activity distinguishes Alzheimer’s disease from healthy aging: Evidence from functional MRI. Proceedings of the National Academy of Sciences of the United States of America, 2004. 101(13): p. 4637–4642.

31. Palmqvist, S., et al., Earliest accumulation of β-amyloid occurs within the default-mode network and concurrently affects brain connectivity. Nature communications, 2017. 8(1): p. 1214–1214.

32. De Havas, J.A., et al., Sleep deprivation reduces default mode network connectivity and anti-correlation during rest and task performance. NeuroImage, 2012. 59(2): p. 1745–1751.

33. Badhwar, A., et al., Resting-state network dysfunction in Alzheimer’s disease: A systematic review and meta-analysis. Alzheimer’s & dementia, 2017. 8: p. 73–85.

34. Sala-Llonch, R., D. Bartrés-Faz, and C. Junqué, Reorganization of brain networks in aging: a review of functional connectivity studies. Front Psychol, 2015. 6: p. 663.

35. Zhou, J., et al., Divergent network connectivity changes in behavioural variant frontotemporal dementia and Alzheimer’s disease. Brain, 2010. 133(5): p. 1352–1367.

36. Ma, N., et al., How Acute Total Sleep Loss Affects the Attending Brain: A Meta-Analysis of Neuroimaging Studies. Sleep, 2015. 38(2): p. 233–240.

37. Zuurbier, L.A., et al., Cerebral small vessel disease is related to disturbed 24-h activity rhythms: a population-based study. European journal of neurology, 2015. 22(11): p. 1482–1487.

38. Mullington, J.M., et al., *Cardiovascular, Inflammatory,* and Metabolic Consequences of Sleep Deprivation. Progress in Cardiovascular Diseases, 2009. 51(4): p. 294–302.

39. Sommer, R., et al., Disrupted Rest-Activity Rhythms and Cerebral Small Vessel Disease Pathology in Older Adults. Stroke, 2021. 52(7): p. 2427–2431.

40. Del Brutto, O.H., et al., Poor sleep quality and silent markers of cerebral small vessel disease: a population-based study in community-dwelling older adults (The Atahualpa Project). Sleep Medicine, 2015. 16(3): p. 428–431.

41. Ning, J., et al., Association of sleep behaviors with white matter hyperintensities and microstructural injury: a cross-sectional and longitudinal analysis of 26,354 participants. Sleep, 2023.

42. Iturria-Medina, Y., et al., Early role of vascular dysregulation on late-onset Alzheimer’s disease based on multifactorial data-driven analysis. Nat Commun, 2016. 7: p. 11934.

43. Thomas, J., et al., Sleep-Cognition Hypothesis In maritime Pilots, what is the effect of long-term work-related poor sleep on cognition and amyloid accumulation in healthy middle-aged maritime pilots: methodology of a case–control study. BMJ Open, 2019. 9(6): p. e026992.

44. Andersson, J.L., S. Skare, and J. Ashburner, How to correct susceptibility distortions in spin-echo echo-planar images: application to diffusion tensor imaging. Neuroimage, 2003. 20(2): p. 870–88.

45. Smith, S.M., et al., Advances in functional and structural MR image analysis and implementation as FSL. Neuroimage, 2004. 23 **Suppl 1**: p. S208–19.

46. Jenkinson, M., et al., Improved optimization for the robust and accurate linear registration and motion correction of brain images. Neuroimage, 2002. 17(2): p. 825–841.

47. Pruim, R.H.R., et al., ICA-AROMA: A robust ICA-based strategy for removing motion artifacts from fMRI data. NeuroImage, 2015. 112: p. 267–277.

48. Fazekas, F., et al., MR signal abnormalities at 1.5 T in Alzheimer’s dementia and normal aging. AJR Am J Roentgenol, 1987. 149(2): p. 351–6.

49. Wardlaw, J.M., et al., Perivascular spaces in the brain: anatomy, physiology and pathology. Nat Rev Neurol, 2020. 16(3): p. 137–153.

50. Doubal, F.N., et al., Enlarged perivascular spaces on MRI are a feature of cerebral small vessel disease. Stroke, 2010. 41(3): p. 450–454.

51. Alsop, D.C., et al., Recommended implementation of arterial spin-labeled perfusion MRI for clinical applications: A consensus of the ISMRM perfusion study group and the European consortium for ASL in dementia. Magn Reson Med, 2015. 73(1): p. 102–16.

52. Andersson, J.L., M. Jenkinson, and S. Smith, Non-linear registration, aka Spatial normalisation FMRIB technical report TR07JA2. FMRIB Analysis Group of the University of Oxford, 2007. 2(1): p. e21.

53. Zhang, Y., M. Brady, and S. Smith, Segmentation of brain MR images through a hidden Markov random field model and the expectation-maximization algorithm. IEEE transactions on medical imaging, 2001. 20(1): p. 45–57.

54. Jenkinson, M., et al., FSL. Neuroimage, 2012. 62(2): p. 782–90.

55. Nickerson, L.D., et al., Using Dual Regression to Investigate Network Shape and Amplitude in Functional Connectivity Analyses. Frontiers in Neuroscience, 2017. 11: p. 115.

56. Beckmann, C.F., et al., Group comparison of resting-state FMRI data using multi-subject ICA and dual regression. Neuroimage, 2009. 47(Suppl 1): p. S148.

57. Goulden, N., et al., The salience network is responsible for switching between the default mode network and the central executive network: Replication from DCM. NeuroImage, 2014. 99: p. 180–190.

58. Li, C., et al., Abnormal Brain Network Connectivity in a Triple-Network Model of Alzheimer’s Disease. Journal of Alzheimer’s Disease, 2019. 69: p. 237–252.

59. Dai, X.-J., et al., Long-term total sleep deprivation decreases the default spontaneous activity and connectivity pattern in healthy male subjects: a resting-state fMRI study. Neuropsychiatric Disease and Treatment, 2015. 11: p. 761–772.

60. Balan, I., et al., Sleep Deprivation in Middle Age May Increase Dementia Risk: A Review. Cureus, 2023. 15(4).

61. Carvalho, C. and P.I. Moreira, Oxidative Stress: A Major Player in Cerebrovascular Alterations Associated to Neurodegenerative Events. Front Physiol, 2018. 9: p. 806.

62. Lloret, A., et al., Is Oxidative Stress the Link Between Cerebral Small Vessel Disease, Sleep Disruption, and Oligodendrocyte Dysfunction in the Onset of Alzheimer’s Disease? Frontiers in Physiology, 2021. 12.

63. Yaffe, K., et al., Sleep Duration and White Matter Quality in Middle-Aged Adults. Sleep, 2016. 39(9): p. 1743–1747.

64. Lutsey, P.L., et al., Sleep apnea, sleep duration and brain MRI markers of cerebral vascular disease and alzheimer’s disease: The Atherosclerosis Risk in Communities Study (ARIC). PloS one, 2016. 11(7): p. e0158758.

65. Lloret, A., et al., Obstructive sleep apnea: arterial oxygen desaturation coincides with increases in systemic oxidative stress markers measured with continuous monitoring. Free radical biology & medicine, 2007. 42(6): p. 893–895.

66. Minakawa, E.N., K. Wada, and Y. Nagai, Sleep Disturbance as a Potential Modifiable Risk Factor for Alzheimer’s Disease. International Journal of Molecular Sciences, 2019. 20(4): p. 803.

67. Carvalho, D.Z., et al., Association of Polysomnographic Sleep Parameters With Neuroimaging Biomarkers of Cerebrovascular Disease in Older Adults With Sleep Apnea. Neurology, 2023. 101(2): p. e125–e136.

68. Daulatzai, M.A., Evidence of neurodegeneration in obstructive sleep apnea: Relationship between obstructive sleep apnea and cognitive dysfunction in the elderly. Journal of Neuroscience Research, 2015. 93(12): p. 1778–1794.

69. Weijs, R.W.J., et al., Longitudinal changes in cerebral blood flow and their relation with cognitive decline in patients with dementia: Current knowledge and future directions. Alzheimer’s & Dementia, 2023. 19(2): p. 532–548.

70. Asllani, I., Effects of 48hr sleep deprivation on cerebral blood flow measured with arterial spin labeling MRI. 2007.

71. Poudel, G.R., C.R.H. Innes, and R.D. Jones, Cerebral Perfusion Differences Between Drowsy and Nondrowsy Individuals After Acute Sleep Restriction. Sleep, 2012. 35(8): p. 1085–1096.

72. Uleman, J.F., et al., Simulating the multicausality of Alzheimer’s disease with system dynamics. Alzheimer’s & Dementia. **n/a**(n/a).

73. López-García, S., et al., Sleep Time Estimated by an Actigraphy Watch Correlates With CSF Tau in Cognitively Unimpaired Elders: The Modulatory Role of APOE. Frontiers in Aging Neuroscience, 2021. 13.

74. Jørgensen, J.T., et al., Shift work and incidence of dementia: A Danish Nurse Cohort study. Alzheimer’s & Dementia, 2020. 16(9): p. 1268–1279.

75. Shi, L., et al., Sleep disturbances increase the risk of dementia: A systematic review and meta-analysis. Sleep Medicine Reviews, 2018. 40: p. 4–16.

